# Nuclei Isolation Protocol for Single-Nucleus RNA Sequencing of Human Stem Cell-Derived Grafts

**DOI:** 10.1101/2025.08.27.672369

**Authors:** Edoardo Sozzi, Petter Storm, Alessandro Fiorenzano

## Abstract

Single-nucleus RNA sequencing enables high-resolution transcriptomic profiling of brain tissue, facilitating detailed analysis of cell identity in models of neurodegeneration and repair. Here, we describe a protocol for isolating nuclei from long-term human stem cell–derived grafts in the rat brain, incorporating vibratome sectioning, graft dissection, nuclear extraction, and fluorescence-activated sorting. This workflow supports analysis of human neurons embedded within host tissue or sensitive to dissociation, offering a powerful approach to assess graft composition, integration, and neuronal identity in living brain.

For complete details on the use and execution of this protocol, please refer to *Fiorenzano et al*.^*1*^

**Subject areas:** Single nucleus RNA sequencing, stem cell biology, neuroscience, transplantation, cell replacement therapy

**Graphical abstract:** 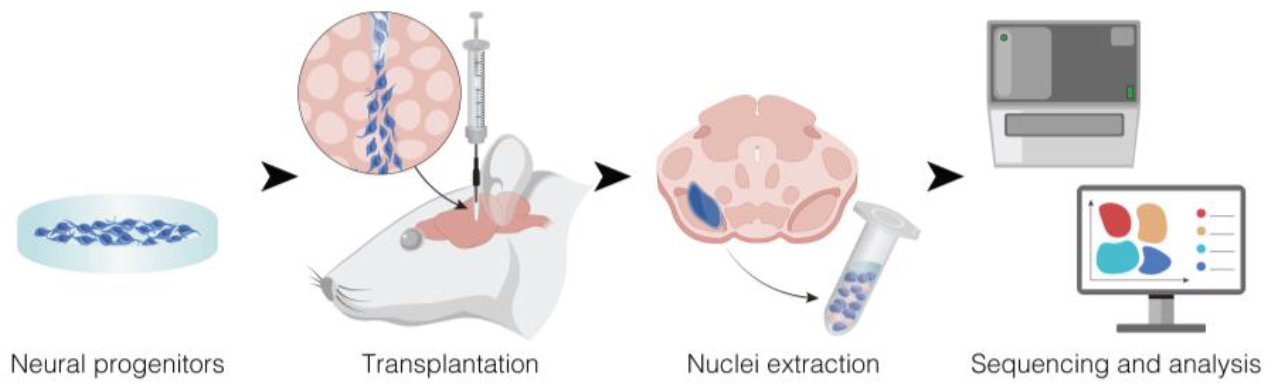

**Highlights:**

- Step-by-step vibratome sectioning of xenografted rat brain tissue.
- Precise dissection of human stem cell–derived grafts from host brain sections.
- Extraction of intact nuclei from fragile, grafted neurons.
- Isolation of single nuclei via fluorescence-activated nuclei sorting (FANS) for snRNA-seq sample preparation.

## Before you begin

The advent of single-cell sequencing has significantly deepened our understanding of cellular diversity in human tissues and has profoundly shaped stem cell biology^2^ - advancing both *in vitro* modeling of human-specific processes and the development of stem cell–based therapies for brain repair^3-7^. However, single-cell RNA sequencing (scRNA-seq) depends on mechanical and enzymatic dissociation, which can compromise cellular integrity, particularly in complex or three-dimensional tissues^8-11^. As a result, fragile cell types are often underrepresented or lost, limiting the generation of accurate and comprehensive cellular maps in complex samples^12,13^.

While scRNA-seq captures both cytoplasmic and nuclear transcripts by profiling whole cells, single-nucleus RNA sequencing (snRNA-seq) focuses on nuclear RNA. Although this can offer a more limited view, it enables analysis of frozen, fixed, or otherwise hard-to-dissociate tissues, reduces dissociation-induced stress and artifacts, and improves recovery of vulnerable and structurally complex cells such as neurons, preserving RNA integrity^14,15^. Thus, snRNA-seq is a powerful alternative for transcriptomic profiling under challenging conditions - such as with primary samples, mature organoid-based models, or xenografts.

Here, we present a detailed protocol for isolating high-quality single nuclei from xenografted brain tissue collected 9–12 months post-transplantation in a rat model of Parkinson’s disease (PD)^1^. The protocol includes step-by-step procedures for vibratome sectioning, graft isolation, nuclear extraction, buffer preparation, and fluorescence-activated nuclei sorting (FANS). We have successfully applied this method across several studies^1,6,16-18^, using stem-cell derived or reprogrammed dopaminergic neuronal grafts across both homotopic and heterotopic transplantation sites. This protocol has also been applied to early-stage grafts (up to 4 weeks post-transplantation), as well as for the isolation of nuclei from ventral midbrain organoids and cell pellets.

These results demonstrate the protocol’s robustness and its broad versatility across different graft types, anatomical locations, and experimental conditions.

For the data presented here, the starting material consists of human ventral midbrain progenitors grafted into a rat model of PD^1,19^. Adult rats were unilaterally lesioned with 6-hydroxydopamine (6-OHDA) injected into the right medial forebrain bundle to induce dopaminergic neuron degeneration. Four weeks after 6-OHDA injection, 150.000 ventral midbrain progenitors were transplanted into the lesioned midbrain, evenly distributed across two stereotaxic injection sites (75.000 cells per site). Tissue was collected 9–12 months post-transplantation for single-nucleus RNA sequencing, following the protocol described here. For detailed lesioning and transplantation procedures, refer to *Fiorenzano et al*.^*1*^

To maximize efficiency and reduce processing time per brain, this protocol is best performed by a team of two to three researchers. Ideally, one researcher should perform surgery and brain extraction, a second prepares vibratome sections, and a third handles graft dissection. If only two researchers are available, one can manage both surgery and graft dissection, while the other focuses on tissue sectioning (Figure 1A). Downstream steps, including nuclei extraction and purification, are typically manageable by a single researcher, depending on the number of samples being processed.

**Figure 1:**
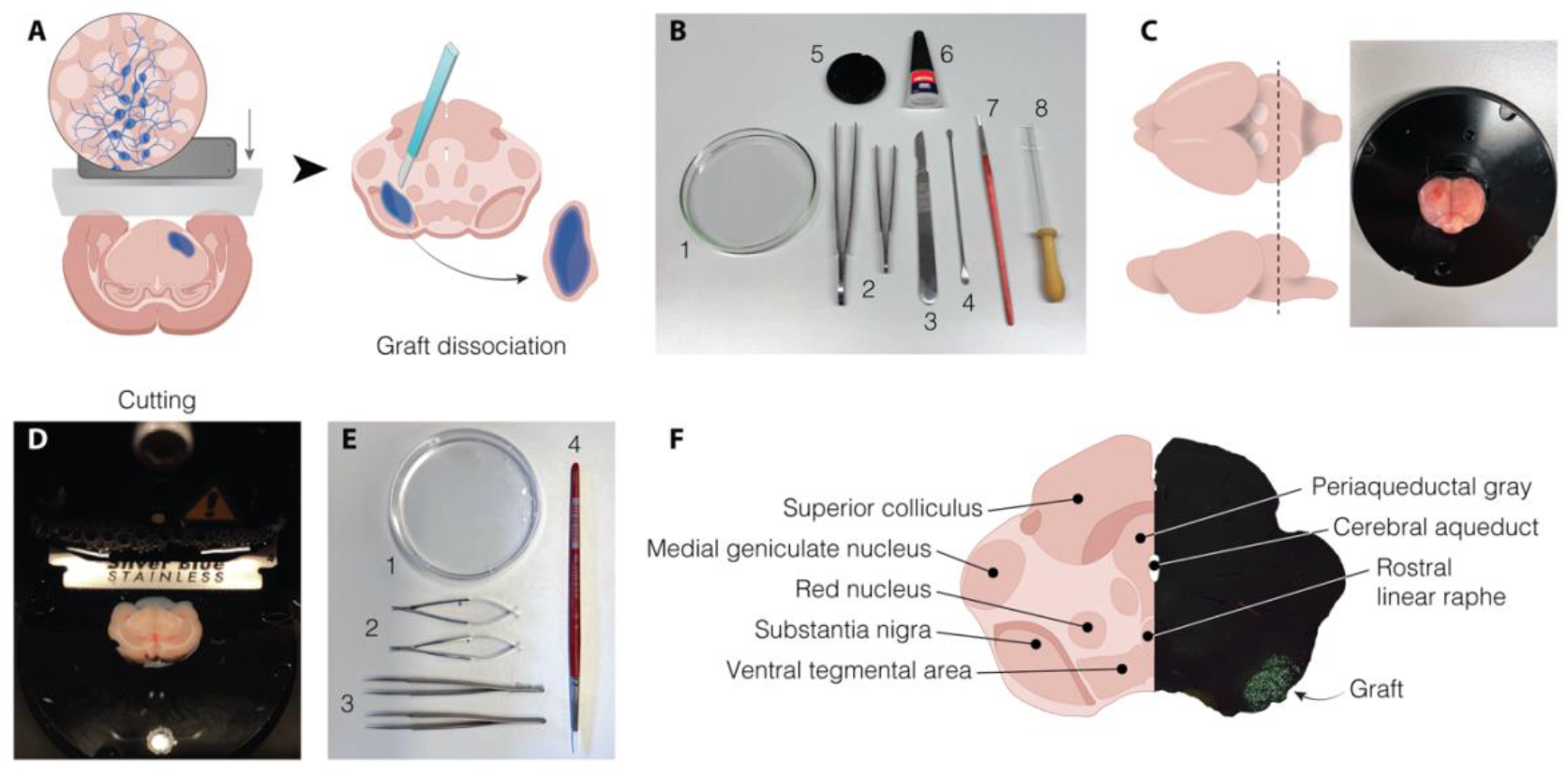
Dissection of human grafts from acute rat brain slices. **A)** Schematic illustration of acute slices prepared from a rat brain and the manual dissection of the graft. **B)** Tools used to prepare for the tissue block for vibratome sectioning. The image shows: (1) glass Petri dish, (2) straight forceps, (3) scalpel, (4) micro spatula, (5) vibratome specimen plate, (6) liquid super glue, (7) soft paintbrush, and (8) glass Pasteur pipette with rubber bulb. **C)** Illustration (left) and picture (right) showing the steps to prepare the tissue block for cutting. **D)** Vibratome sectioning of the rat brain submerged in aCSF. **E)** Instruments used for graft dissection. The image shows: (1) plastic Petri dish for collecting slices, (2) spring scissors, (3) straight forceps, and (4) fine-tipped paintbrush for tissue collection. **F)** Schematic overview of the main anatomical nuclei in the midbrain to be used as a reference for graft dissection, and to assess the correct placement of the graft.

### Institutional permissions

All animal procedures were conducted in accordance with the European Union Directive 2010/63/EU and were approved by the local ethical committee at Lund University and the Swedish Department of Agriculture (*Jordbruksverket*). Female athymic nude rats were obtained from Envigo and housed in ventilated cages with *ad libitum* access to food and water, under a 12-hour light/dark cycle. The strain and sex of the animals are not critical for the execution of this protocol. Researchers must obtain all necessary institutional and governmental approvals before performing animal experiments.

### Key resources table

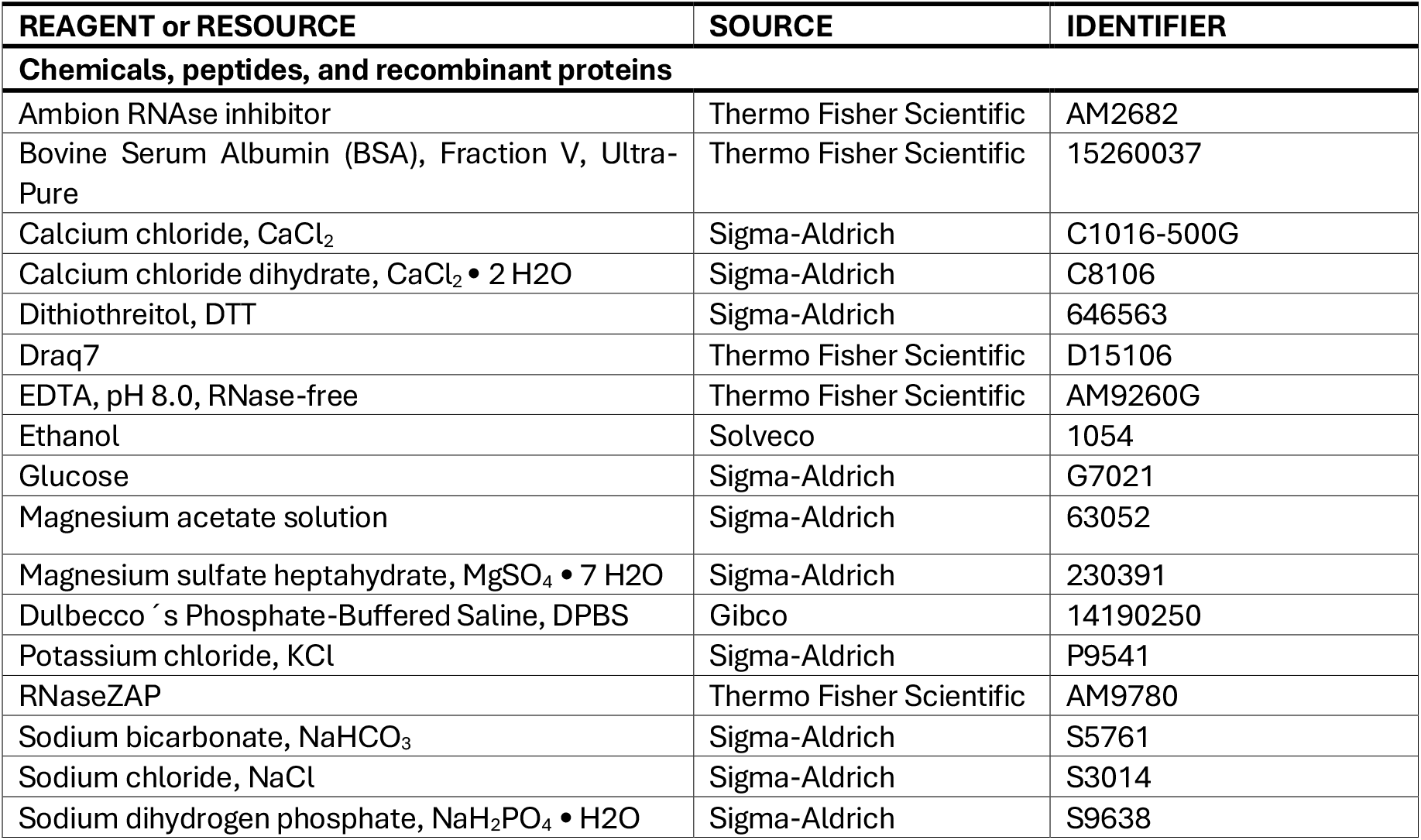
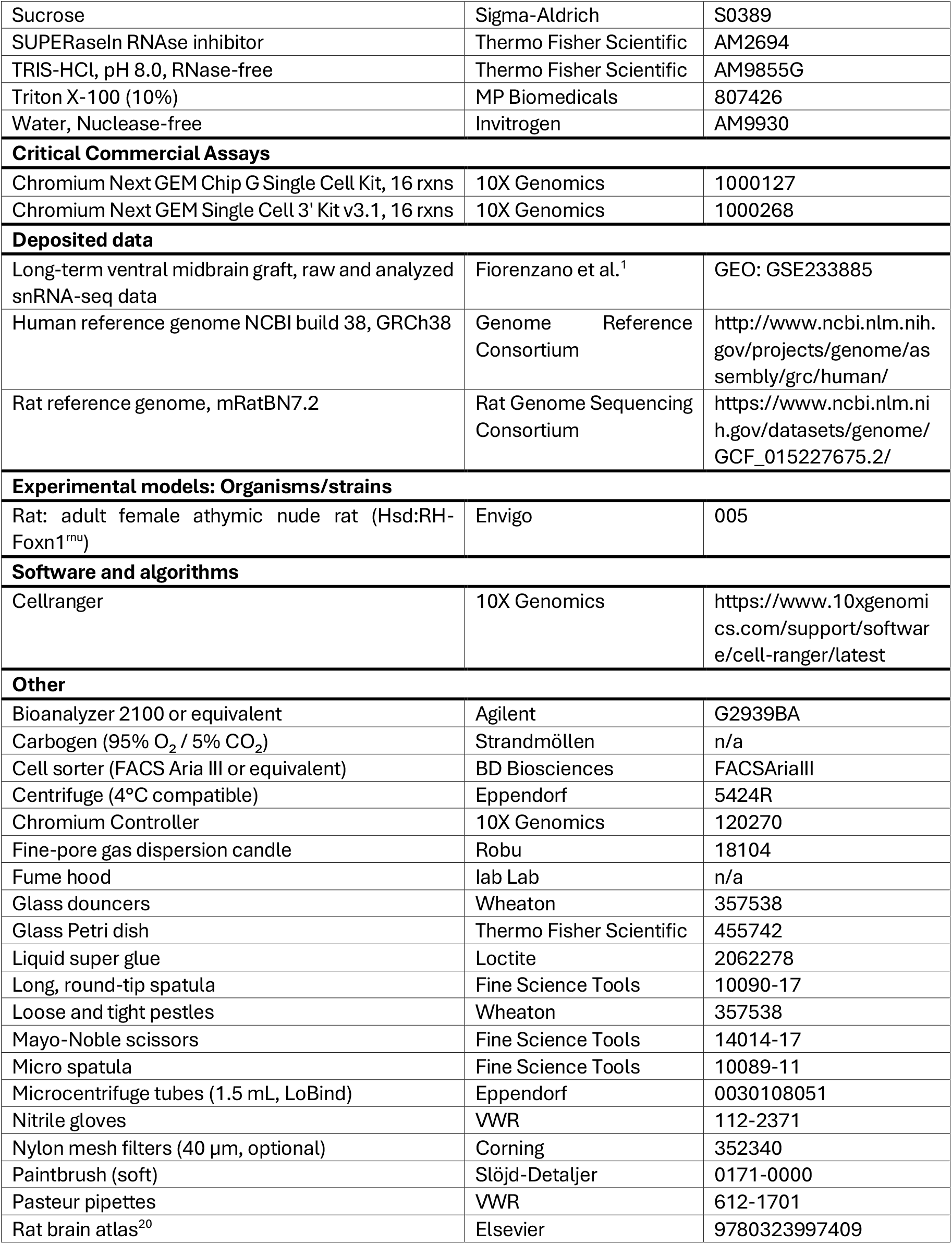
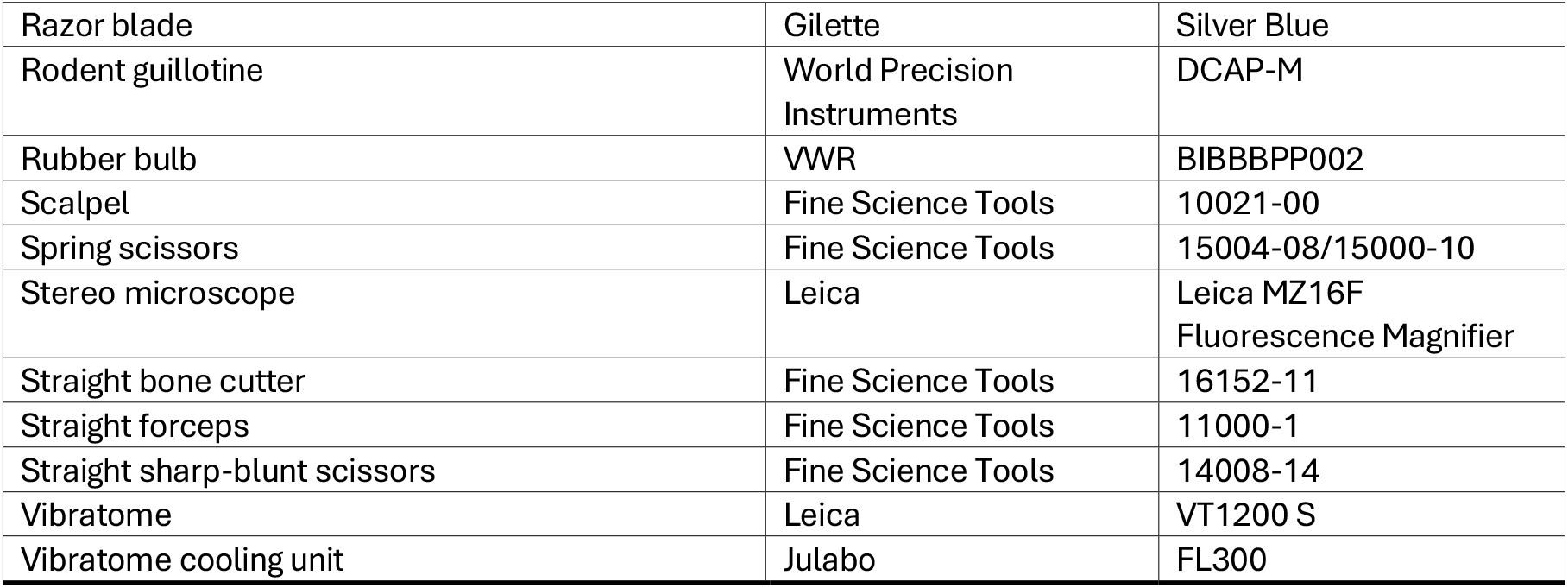

## Materials and equipment setup

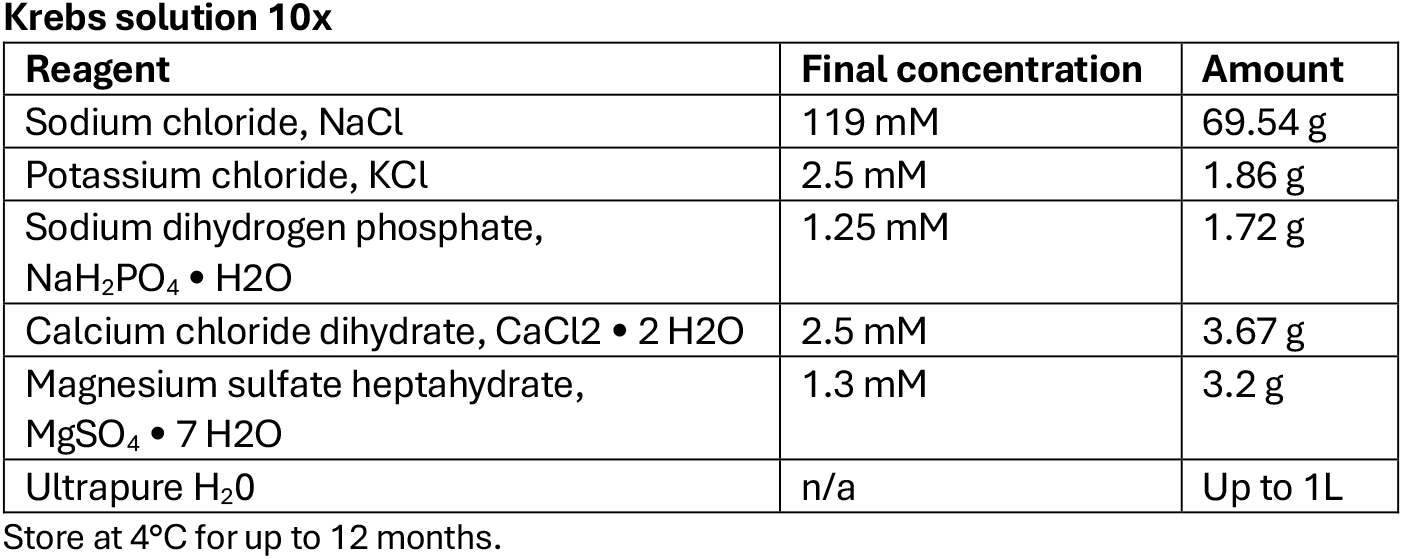

In an 1L volumetric flask add 100 ml distilled water, then add the listed reagents under constant magnetic stirring; when all reagents have been added use distilled water to reach the volume of 1L.

Artificial cerebrospinal fluid solution (aCSF – Ringer solution)
| Reagent | Final concentration | Amount |
| --- | --- | --- |
| Sodium bicarbonate, NaHCO<sub>3</sub> | 26 mM | 2.18 g |
| Glucose | 25 mM | 4.95 g |
| Krebs solution (10x) | n/a | 100 mL |
| Ultrapure H<sub>2</sub>O | n/a | Up to 1L |

Store at 4°C for up to one month.

In an 1L volumetric flask add 100 ml distilled water, add the listed reagents under constant magnetic stirring. When all reagents have been included, add 100 ml of 10x Krebs solution and distilled water to reach the volume of 1L. Leave the solution bubbling with a 95% O_2_ 5% CO_2_ gas mixture (Carbogen) for one hour. The final solution should have an osmolarity ranging between 290-310 mOsm/L and a pH around 7.4.

Lysis buffer for nuclei extraction
| Reagent | Final concentration | Amount |
| --- | --- | --- |
| Nuclease-free water | n/a | 11.4 mL |
| Sucrose (1.5 M) | 0.32 M | 3.2 mL |
| Calcium chloride, CaCl<sub>2</sub> (1 M) | 5 mM | 75 µL |
| Magnesium acetate solution (1 M) | 3 mM | 45 µL |
| EDTA (0.5 M) | 0.1 mM | 3 µL |
| TRIS-HCl, pH 8.0 (1M) | 10 mM | 150 µL |
| *Dithiothreitol, DTT (1M) | 1 mM | 15 µL |
| *Triton X-100 (10%) | 0.1 % | 150 µL |
| *Ambion RNase inhibitor (40 U/µL) | 0.04 U/µL | 15 µL |
| *SUPERaseIn RNase inhibitor (40 U/µL) | 0.04 U/µL | 15 µL |
| **Total** | **n/a** | **15 mL** |

Store at 4°C for up to one month.
\*Component to be added fresh before use. Keep the working solution in ice while in use.

***CRITICAL:*** *Triton X-100 is an irritant. It can cause skin and eye irritation and may be harmful if inhaled or swallowed. Handle in a well-ventilated area or under a fume hood using gloves and safety goggles*.

***CRITICAL:*** *Dithiothreitol is toxic and a strong reducing agent. It can cause respiratory and skin irritation and is harmful if swallowed or inhaled. Always use in a fume hood to avoid inhalation and handle with gloves and eye protection*.

Resuspension buffer for nuclei extraction
| Reagent | Final concentration | Amount |
| --- | --- | --- |
| RNase-free DPBS (1X) | n/a | 4.93 mL |
| Bovine Serum Albumin, Fraction V, Ultra-Pure (7.5 %) | 0.1 % | 66 µL |
| *Ambion RNase inhibitor (40 U/µL) | 0.04 U/µL | 5 µL |
| *SUPERaseIn RNase inhibitor (40 U/µL) | 0.04 U/µL | 5 µL |
| **Total** | **n/a** | **5 mL** |

Store at 4°C for up to one month.
\*Component to be added fresh before use. Keep the working solution in ice while in use.

## Step-by-step method details

### Brain tissue sectioning

**Timing: 30 min for preparations; 1 h per brain processed**.

*This step involves the preparation of acute brain slices from freshly extracted rat brains serving as the starting material for subsequent graft dissection and nuclei isolation (Figure 1A). Preserving tissue viability and structural integrity is critical throughout the process*.

***Note***: *Autoclave all surgical, dissection, and cutting tools before use*.

***CRITICAL:*** *All animal procedures must follow institutional and national guidelines for ethical animal research*.

***CRITICAL:*** *Wear safety goggles to protect against bone fragments during skull removal and dissection. Always use protective clothing, gloves, and a face mask throughout the procedure*.

***CRITICAL:*** *All steps must be performed under PFA-free conditions. Benches, tools, and solutions should never be shared with PFA-treated samples, as even trace contamination can compromise RNA integrity, and make tissue unsuitable for nuclei isolation and RNA sequencing*.

1. Setting up the vibratome and the cutting station.
  a. Prepare 500-700mL of ice-cold artificial cerebrospinal fluid (aCSF) and bubble it continuously with carbogen gas (95% O_2_ / 5% CO_2_) through a fine-pore candle, or equivalent. Ensure the aCSF is bubbled for at least 30 minutes prior to slicing to achieve proper oxygenation. **Note**: The required volume of aCSF depends on the vibratome model and the number of samples processed. For reference, 500 mL is sufficient when processing 3 samples on a Leica VT1200/S.
  b. Thoroughly clean the vibratome and surrounding workspace with 70% ethanol to maintain sterility and prevent sample contamination. If salt deposits are present on the blade holder or tray, rinse them with water before disinfecting with 70% ethanol.
  c. Turn on the vibratome’s cooling unit at least 15 minutes before use and allow it to reach -4°C. If a different cooling method is preferred, make sure the vibratome’s tray reaches a temperature of -4°C before starting the cutting.
  d. Carefully insert a fresh razor blade, ensuring it is securely fixed, and turn the vibratome on.
  e. Use the VibroCheck tool to verify the blade vibration (acceptable value: |∑µm| < 0.2), adjusting the blade positioning if needed. **CRITICAL**: Improper blade alignment can increase vibration, compromising section quality and integrity for the following steps. Always confirm optimal settings using the VibroCheck before proceeding (see Troubleshooting – Problem 1).
  f. Install the buffer tray onto the vibratome holder and fill it with 300 mL of ice-cold, oxygenated aCSF. Maintain continuous, low intensity bubbling of the aCSF in the tray throughout the cutting using a dispersion stone, without creating turbulence.
  g. Prepare additional instruments and materials next to the vibratome to facilitate efficient workflow during tissue sectioning (Figure 1B). In particular, prepare a glass petri dish (1), straight forceps (2), a sterile scalpel (3), a micro spatula (4), the vibratome specimen plate (5), liquid super glue (6), a soft paintbrush (7) and a glass Pasteur pipette with rubber bulb (8).
2. Preparation of the surgical station.
  a. Prepare all sterile surgical tools on a sterile surface or surgical tray, including sharp-blunt straight scissors, Mayo-Noble scissors, a straight bone cutter, a long, round-tip spatula, and forceps^21^.
  b. Transfer 40mL of pre-chilled, carbogen-saturated aCSF (from step 1a) into a 50mL Falcon tube. Immediately proceed to the following step.
3. Surgery and brain extraction.
  a. Euthanize the animal via rapid decapitation using a small animal decapitator guillotine and place the head on a clean dissection surface.
  b. Use a straight scissor to cut the scalp from the midline from backward forward. Then expose the skull by retracting both sides of the scalp.
  c. Remove any remaining muscle attached to the posterior and inferior skull with a bone cutter.
  d. Cautiously, insert sharp-blunt straight scissors through the foramen magnum and make bilateral incisions (∼1 cm) along the basioccipital bone, exposing the brain stem and cerebellum.
  e. Cut dorsally along the midline, exposing temporal and parietal bones.
  f. Insert the tip of the bone cutter between the lateral side of the brain and the temporal bones and remove them. Make shallow cuts to avoid damaging the underlying brain.
  g. Gently pull up the parietal bones up from the back up and remove them.
  h. Carefully trim the frontal bones using a bone cutter and Mayo-Noble scissors until the olfactory bulbs are exposed. Move back to the inferior surface of the brain to cut trigmeminal and optic nerves.
  i. Using a loung, round-tip spatula, carefully dissect the olfactory bulbs to release and extract the brain.
  j. Rinse the exposed brain with 10 mL of ice-cold, carbogen-saturated aCSF.
  k. Use a long, round-tip spatula to gently scoop out the intact brain and transfer it into the 50mL Falcon tube with the remaining ice-cold aCSF. Do not manipulate the brain directly to prevent tissue damage.
4. Brain sectioning.
  a. Gently transfer the brain from the 50 mL Falcon tube into the glass Petri dish by carefully tilting the tube. Use forceps as needed to guide the tissue, transferring it with minimal amount of aCSF.
  b. Stabilize the brain in the Petri dish using forceps, then perform a clean coronal cut at the level of the cerebellum to create a flat surface for mounting the tissue block (Figure 1C). **Note:** The cutting angle and site may be adapted depending on the area of the graft. The preparation illustrated here facilitates access to both the midbrain and striatum, common grafting targets in PD animal models.
  c. Apply a small amount of cyanoacrylate glue to the specimen disc, spreading it to create an even layer.
  d. Using a microspatula and forceps, slide the brain onto the glued specimen disc, securing the flat cut surface vertically to the vibratome specimen plate (Figure 1C).
  e. Allow approximately 10 s for the glue to solidify and apply a drop of aCSF on the brain to remove excess glue.
  f. Insert the disc into the buffer tray filled with ice-cold, carbogen-satured aCSF (prepared in step 1f). Remove any remaining meninges with fine forceps to prevent interference with the slicing. **CRITICAL:** Complete steps 4c-f within 30 s to preserve tissue viability and slicing quality.
  g. On the vibratome control panel, set the sectioning parameters. The recommended settings used in this protocol are: speed - 0.1 mm/s; amplitude – 1.7 mm; slice thickness 275 μm. **Note:** Parameters may require adjustment based on graft location, size, and time after transplantation. For fragile or mature grafts, reduce speed to minimize tissue damage and preserve structural integrity. Generally, suggested ranges are: speed 0.05-0.2 mm/s; amplitude 1.0-1.7 mm; slice thickness 250-400 μm (see Troubleshooting – Problem 1).
  h. Begin sectioning from the ventral side and proceed dorsally (Figure 1D). **Note:** Ensure the blade remains fully submerged in chilled aCSF throughout the procedure.
  i. Use a soft paintbrush and a glass Pasteur pipette to gently handle and transfer slices for downstream processing.

### Graft dissection and isolation

**Timing: 10 min for preparation; 45 min per graft processed**.

*This step involves isolating the graft region from acute brain slices for downstream nuclei extraction and molecular analyses (Figure 1A)*.

***CRITICAL:*** *Autoclave all dissection tools prior to use and keep them free from RNAse contamination. Complete dissection within 1 hour from sectioning to preserve RNA integrity* (see Troubleshooting – Problem 2).

***CRITICAL:*** *Always wear gloves and a face mask during dissection*.

5. Preparation of the dissection station.
  a. Wipe down all work surfaces and the dissection microscope with 70% ethanol and RNaseZAP.
  b. Turn on the dissection microscope and adjust the illumination as needed.
  c. Set up all necessary instruments next to the microscope (Figure 1E). In particular, prepare a sterile glass or plastic Petri dish for collecting slices (1), spring scissors (2), straight forceps (3), and a fine-tipped paintbrush (4). Also prepare a box of dry ice and pre-labeled 1.5 mL microcentrifuge tubes (one per dissected graft slice, see below).
6. Graft dissection
  a. Transfer the brain slices from step 4f into the Petri dish together with chilled, carbogen-saturated aCSF. Ensure slices are fully submerged.
  b. Under the dissection microscope, use the forceps and paintbrush to carefully unfold and examine each slice, identifying those containing the graft. In the experimental setup described here, the graft is typically localized within 2-4 consecutive slices per animal (see Troubleshooting – Problem 3). **Note:** if grafted cells express a fluorescent marker (e.g., GFP), use fluorescence to aid the identification (Figure 1F, right). Otherwise, rely on anatomical landmarks and a rat brain atlas^20^ to locate the area to dissect (Figure 1F, left).
  c. Use the forceps to stabilize the slice. Then, carefully cut around the graft using spring scissors, without disrupting the surrounding tissue.
  d. Use the paintbrush (Figure 1E, 4) or straight forceps (3) to transfer each dissected graft into a separate 1.5 mL microcentrifuge tube, minimizing the amount of aCSF carried over. Immediately place tubes in dry ice. Repeat steps 6c/6d for all the brain slices.
  e. After all the graft pieces are collected, transfer the tubes to a -80°C freezer for long-term storage.

**Pause Point:** Dissected grafts can be stored at –80°C for up to 2 years with minimal RNA degradation, provided they were snap-frozen and kept cold. Avoid freeze-thaw cycles, which can compromise RNA integrity.

### Nuclei extraction

**Timing: 1.5 h (5-10 samples), 2 h (11-15 samples)**.

*This step involves the dissociation of frozen graft tissue (or other sample types) into a nuclear lysate optimized for downstream isolation and sorting (Figure 2A). The protocol is also compatible with cell pellets and snap-frozen organoids*.

**Figure 2:**
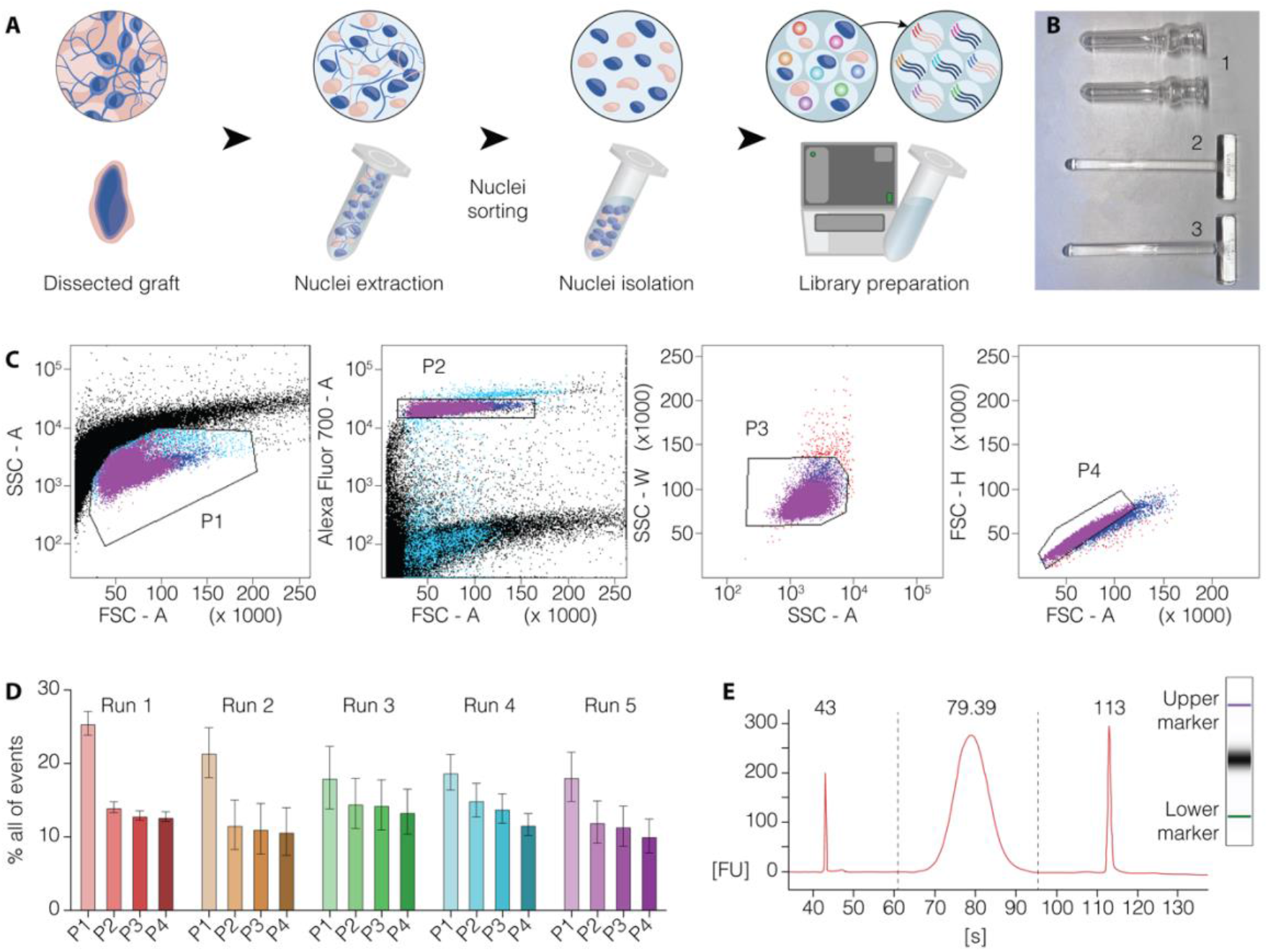
From graft dissection to nuclei isolation, purification, and snRNA-seq library preparation. **A)** Schematic overview of the workflow from dissected graft tissue through nuclei extraction, purification by FANS, and library preparation for snRNA-seq. **B)** Image of the douncers (1) used for mechanical dissociation, along with the loose-fitting (2) and tight-fitting pestles (3). **C)** Representative gating strategy for the isolation of intact single nuclei for downstream snRNA-seq. **D)** Barplots illustrating the proportion of events in each gate across 5 representative sorting runs (n=3-5). **E)** Electropherogram trace of a high-quality snRNA-seq library, highlighting the upper and lower size markers and a central peak corresponding to the expected library fragment size.

***Note:*** *It is not recommended to process more than 15 samples per run, as prolonged thawing or processing time increases the risk of RNA degradation* (see Troubleshooting – Problem 2).

***CRITICAL:*** *Work in an RNase-free environment to reduce the risk of RNA degradation. Always wear gloves and protective clothes during this procedure*.

***CRITICAL:*** *Temperature control is essential throughout nuclei extraction. Perform all steps on ice or at 4°C to prevent RNA degradation*.

7. Set up the nuclei extraction station
  a. Clean the entire bench surface, pipettes, and any tube racks thoroughly with RNaseZAP.
  b. Place glass douncers on regular ice to chill (Figure 2B).
  c. Cool the centrifuge to 4°C and ensure all centrifuge-compatible adaptors are pre-cooled.
  d. Place two pre-labeled DNA LoBind 1.5 mL tubes per sample on ice. Include at least one extra tube for FANS reanalysis (see step 11).
  e. Prepare lysis (1 mL/sample) and resuspension (600 µL/sample) buffers on ice, complementing the base lysis solution with DTT, Triton X-100, and RNAse inhibitors.

**CRITICAL:** Always handle Triton X-100 and DTT under a fume hood with appropriate personal protection to avoid respiratory irritation or other health hazards. See *the Material and Equipment Setup* section for additional information.

**Note:** It is recommended to prepare additional buffer volume in case of large or hard-to-dissociate grafts. For 8 mature graft samples (e.g. 9-12 months post-transplantation), 10 mL of lysis buffer and 5 mL of resuspension buffer are typically sufficient (see Troubleshooting – Problem 4).

8. Tissue processing
  a. Thaw dissected grafts on ice for 5 minutes.
  b. Add 500 µL of cold lysis buffer directly into each tube containing a graft. Gently pipette up and down using a P1000 to detach the tissue from the tube walls.
  c. Transfer the tissue into a pre-chilled glass douncer using the same P1000 tip. Also transfer any remaining lysis buffer from the tube, including visible debris.
  d. Add an additional volume of 500 µL of cold lysis buffer to the douncer. **Note:** If debris still remains in the original tube, use this second 500 µL to rinse and transfer it into the douncer.
  e. Homogenize the sample using a loose-fitting pestle (10-15 strokes), then switch to a tight-fitting pestle (10-15 additional strokes) until the solution becomes uniformly cloudy and small bubbles are visible (see Troubleshooting – Problem 4).
  f. Transfer the homogenized lysate into a cold 1.5 mL DNA LoBind tube (from step 7d). No large tissue fragments should be visible.
  g. Centrifuge the lysate at 900 g for 15 minutes at 4°C. Ensure the centrifuge is balanced and kept cold throughout the spin.
9. Coating and preparation for FANS
  a. Add 100 µL of resuspension buffer to the designated reanalysis tube, on ice.
  b. Add Draq7 at 1:1000 dilution to the remaining resuspension buffer. Mix gently and keep cold. **Note:** Draq7 should only be added to the buffer intended for FANS collection and not to the reanalysis tube.
  c. Coat the cold DNA LoBind collection tubes (prepared in step 7d) by adding up to 200 µL of Draq7-containing resuspension buffer to each tube. Keep them on ice. **CRITICAL:** Coating should be performed at least 30 minutes before sorting to minimize nuclei adherence to tube walls (see Troubleshooting – Problem 5).
10. Pellet resuspension
  a. After centrifugation (step 8g), carefully aspirate and discard the supernatant using a P200 pipette, without disrupting the pellet. The supernatant should appear clear or slightly cloudy but without visible tissue clamps.
  b. Close the tube and gently flick the bottom to help detach the pellet from the walls.
  c. Add 200–400 µL of resuspension buffer using a P200 pipette, adjusting the volume based on graft size. Keep the tubes in ice and immediately proceed to the isolation step via FANS.

**Note:** For large grafts (e.g., 6 months post-transplantation), 300 µL of resuspension buffer generally yields an optimal nuclear concentration while minimizing aggregates that could clog the sorter; prepare additional buffer in advance in case dilution is needed during sorting. If the buffer used for tube coating (step 9c) has incubated for at least 30 minutes, it can also be removed and reused for this step (see Troubleshooting – Problem 6).

**Optional**: When working with cell or organoid pellets, filter the resuspended nuclei through a 40 µm nylon mesh to remove large debris and minimize the risk of clogs during FANS. Note that this step is not always recommended for graft samples, as it may lead to significant material loss (see Troubleshooting – Problem 6).

### Nuclei purification via Fluorescence-Activated Nuclei Sorting (FANS) and snRNA-seq library preparation

**Timing: sorting, 2 h (5-10 samples), 3 h (11-15 samples); library generation, 5h**.

*This step involves the isolation of intact single nuclei from debris and doublets using fluorescence-activated nuclear sorting (FANS) based on size and DNA content (Draq7). An accurate gating strategy is essential to ensure a high-purity nuclear population for downstream single-nucleus RNA sequencing*.

11 Nuclei purification via FANS.
  a. Prepare the cell sorter with a 100 µm nozzle and activate the red laser (640 nm) for Draq7 detection.
  b. Set up the gating strategy by first excluding debris (P1, Figure 2C) and selecting the population of interest based on size and Draq7 staining (P2, Figure 2C). Then remove doublets by plotting side scatter width (SSC-W) vs side scatter area (SSC-A) (P3, Figure 2C), followed by forward scatter height (FSC-H) vs forward scatter area (FSC-A) (P4, Figure 2C). **Note:** When running this protocol for the first time, include an unstained control sample to establish the gating strategy. For subsequent runs, this step is optional but recommended, as it ensures consistency between distinct runs (Figure 2C,D).
  c. Remove any residual coating solution from the reanalysis tube and insert it in the collection position of the sorter.
  d. Sort ≈200-500 nuclei into a reanalysis tube to verify that the sort is proceeding correctly, adjusting gates as necessary. Reanalyze the sorted nuclei to confirm the purity of the gated population. It is recommended to perform this quality control step on the most abundant sample (see Troubleshooting – Problem 7).
  e. Sort each sample into a separate, pre-coated collection tube. Remove any residual coating solution before sorting. Keep all tubes on ice after sorting and proceed immediately to the library preparation step.
12 Preparation of 10X Genomics libraries for snRNA-seq
  a. Centrifuge the nuclei suspension at 500 g for 10 min at 4°C. Carefully remove the supernatant without disturbing any visible pellet. Retain a final volume of 34 µL for loading.
  b. Thaw all 10X reagents on ice.
  c. Load the nuclei suspension and 10X reagents into a Chromium Chip according to the manufacturer’s instructions. Proceed with GEM generation and library preparation using the 10X Genomics Single Cell 3’ (v3.1 or later) protocol adapted for nuclei. **CRITICAL: Always use freshly prepared nuclei**, as prolonged delays can compromise RNA integrity and significantly diminish library quality and sequencing outcomes.

f. Verify the size distribution and integrity of the prepared libraries by loading 1 µL on an Agilent Bioanalyzer. A representative high-quality trace is presented in Figure 2E (see also Troubleshooting – Problem 2).

### Expected outcomes

This protocol enables the reproducible isolation of high-quality, intact single nuclei from human stem cell–derived grafts dissected from acute rat brain slices. Following FANS, users can expect to recover approximately 20.000 to 100.000 nuclei per graft, depending on graft size and maturation stage. The sorted nuclei display intact nuclear membranes (indicated by Draq7 exclusion), minimal debris contamination, and well-defined forward and side scatter profiles (Figure 2C,D). These purified nuclei are well-suited for downstream single-nucleus RNA sequencing (snRNA-seq), consistently yielding high-quality cDNA libraries characterized by a dominant peak of 400-500 basepairs, as confirmed by quality control assays such as the Bioanalyzer (Figure 2E).

During data pre-processing, sequencing reads are aligned to a combined human (GRCh38) and rat (mRatBN7.2) reference genome using the 10X Genomics Cell Ranger pipeline, enabling the discrimination between grafted human cells and contaminating host rat cells via the GEM (Gel Bead-in-Emulsion) classification output (see Troubleshooting – Problem 3). In the representative setup described here, up to 40% of the recovered nuclei are of human origin (Figure 3A), with yield varying based on graft size and whether the dissection is guided by GFP fluorescence. This dual-genome alignment approach enables parallel transcriptomic profiling of xenografted human cells and the surrounding rat microenvironment, allowing the investigation of host–graft cellular interactions.

**Figure 3:**
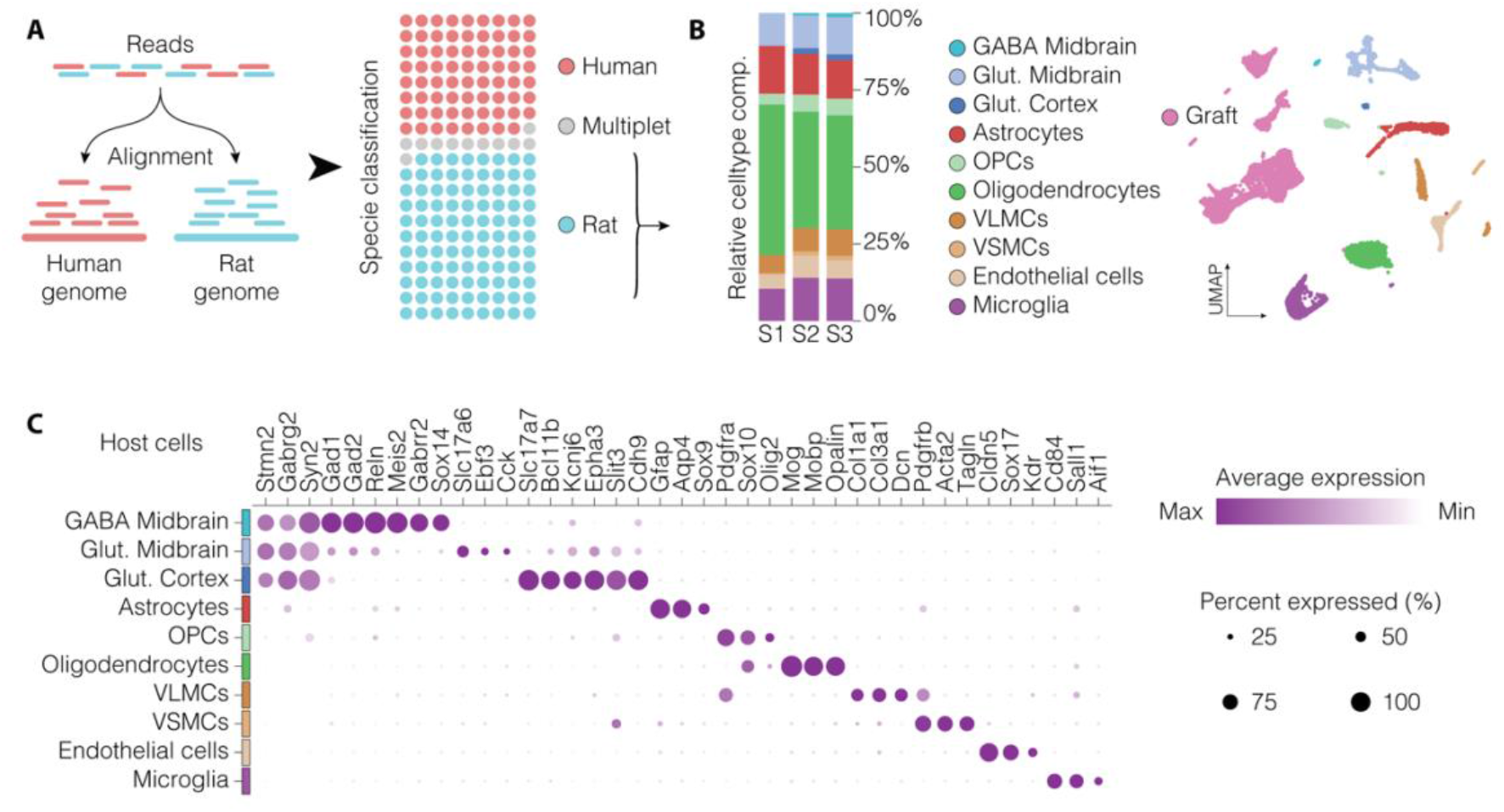
Representative analysis of host cell populations in the graft microenvironment. **A)** Alignment of snRNA-seq reads to a combined human and rat reference genome, enables accurate species discrimination, resulting in an average of ∼40% human nuclei across samples. The use of the cellranger GEM classification method reveals a low proportion of multiplet mismatches, indicating high discriminating specificity. **B)** UMAP visualization (right) and relative quantification (left) of host-derived cell types in the graft microenvironment. **C)** Dot plot showing the expression of canonical marker genes for glutamatergic and GABAergic neurons, astrocytes, oligodendrocytes, perivascular cells, and microglia across identified clusters. OPCs, oligodendrocyte precursor cells; VLMCs, vascular leptomeningeal cells; VSMCs, vascular smooth muscle cells.

Typical 10X Genomics sequencing runs yield approximately 50,000 – 100,000 reads per nucleus with <15% being unmapped and >70% reads that map to barcodes assigned to real cells (vs. background). This level of coverage provides robust gene expression profiling, allowing for detailed molecular characterization and accurate cell type identification of grafted neurons.

## Quantification and statistical analysis

For downstream analysis, sequencing reads are aligned to a combined human (GRCh38) and rat (mRatBN7.2) reference genome, allowing clear separation of human graft-derived nuclei from host rat nuclei for independent characterization (Figure 3A).

In the context of homotopic transplantation into the ventral midbrain, the host-derived nuclei population primarily comprises astrocytes (Gfap, Aqp4, Sox9), oligodendrocytes (Mog, Mobp, Opalin), and oligodendrocyte precursors cells (Sox10, Pdgfra), reflecting the glial response typical of the 6-OHDA lesion model of PD (Figure 3B,C). Additionally, glutamatergic neurons (Slc17a7/vGlut1, Slc17a6/vGlut2, Bcl11b/Ctip2), GABAergic neurons (Sox14, Gabrr2, Gad1, Gad2), perivascular cells (Col1a1, Col3a1, Pdgfrb, Acta2) and microglia (Cd84, Sall1, Aif1) are also reliably detected within host-derived transcriptomic clusters (Figure 3B,C).

For the xenografted human nuclei, stringent quality control (QC) is critical to exclude low-quality nuclei and potential doublets. QC filtering typically involves setting thresholds based on the number of detected features (genes) per nucleus and mitochondrial read content. Notably, data generated using this protocol consistently show mitochondrial read fractions below 1%, indicating high nuclear integrity and low environmental RNA contamination (Figure 4A,B).

**Figure 4:**
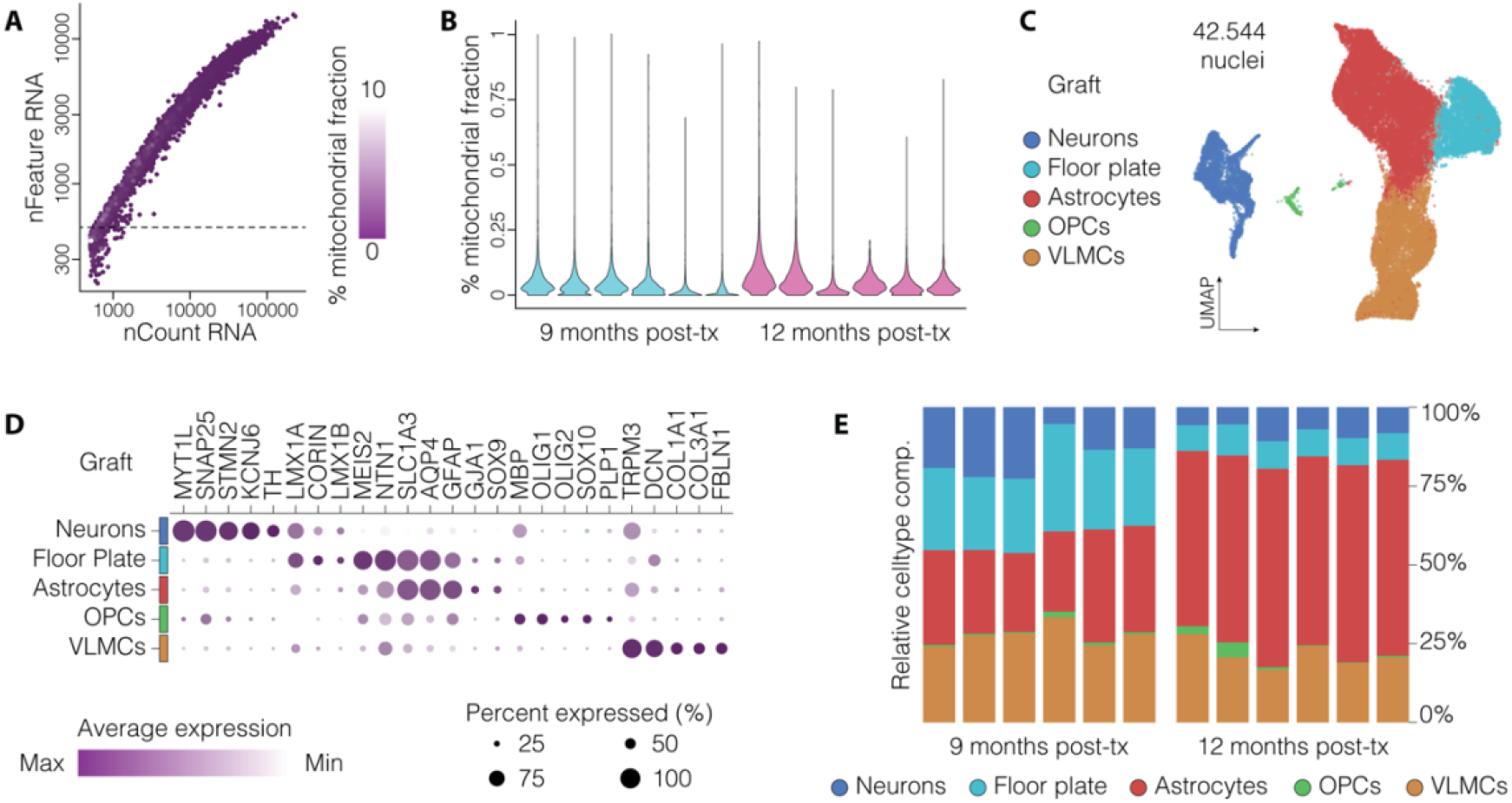
Quality control and representative analysis of human cell types in long-term grafts. **A)** Scatter plot of detected genes (nFeature_RNA) versus total UMI counts (nCount_RNA) per nucleus, colored by percentage of mitochondrial reads. The dotted line marks the gene threshold used to exclude low-quality nuclei from downstream analysis. **B)** Violin plots showing mitochondrial read percentages in 9- and 12-month-old grafts, consistently below 0.3%. **C)** UMAP plot displaying human cell types identified in long-term ventral midbrain grafts, including dopamine neurons, floor plate-like cells, astrocytes, oligodendrocyte precursor cells (OPCs), and vascular leptomeningeal cells (VLMCs). **D)** Dot plot illustrating the expression of selected marker genes across identified cell types. **E)** Bar plots quantifying cell type composition in 9- and 12-month-old grafts, showing consistent profiles across samples.

Other typical downstream analytical steps include:

1. Data integration across graft samples using the Harmony algorithm^22^ to correct for batch effects and other technical artifacts (see Troubleshooting – Problem 8).
2. Clustering via graph-based methods, with cluster structures visualized through Uniform Manifold Approximation and Projection (UMAP) embeddings.
3. Cell type annotation based on identification of differentially expressed genes (DEGs) between clusters, automated cell type annotation in parallel with manual refinement using canonical marker genes.

In the representative example illustrated here, the graft clusters include ventral midbrain (vMB) floor plate–like cells (LMX1A, LMX1B, CORIN), postmitotic dopaminergic neurons (TH, KCNJ6/GIRK2, MYT1L), astrocytes (AQP4, GFAP, GJA1,SOX9), oligodendrocyte progenitor cells (OLIG1/2, SOX10), and vascular leptomeningeal cells (COL1A1, DCN, FBLN1) (Figure 4C,D). Notably, this optimized protocol for isolating human nuclei from long-term grafts resulted in low inter-sample variability and highly reproducible cell clustering across grafts analyzed at 9- and 12-months post-transplantation (Figure 4E).

## Limitations

Single-nucleus RNA sequencing (snRNA-seq) enables transcriptomic profiling from both fresh and frozen tissue, making it particularly well-suited for studying fragile or archived biological samples. However, this methodology presents several limitations that must be considered during experimental design and data interpretation. First, because snRNA-seq captures only nuclear RNA, lower total UMIs/genes detected are expected. Also, the enrichment for unspliced pre-mRNA alters the natural ratio of spliced to unspliced transcripts, potentially compromising advanced bioinformatic analyses relying on transcriptional dynamics, such as RNA velocity. Second, spatial context is lost during mechanical dissociation and nuclear isolation, limiting the ability to associate transcriptional signatures with specific graft regions or anatomical structures. Additionally, migratory cell populations that have left the original graft site may be underrepresented or missed entirely. This limitation can be partially mitigated by labeling grafted cells with a nuclear fluorescent marker (e.g., GFP), allowing for specific sorting during the purification step. Finally, because the entire sample is consumed during nuclei extraction, no material remains from the same graft/animal for follow-up validation experiments such as immunofluorescence staining, or proteomic profiling. These limitations highlight the importance of integrating snRNA-seq with complementary techniques to achieve a more complete understanding of cellular diversity in stem cell-derived grafts.

## Troubleshooting

### Problem 1

Uneven cuts during vibratome sectioning can damage the rat brain, compromising tissue integrity and downstream analyses. This issue commonly arises when the the blade vibrates too much, is cracked or is not properly secured, all of which prevent clean and consistent sectioning.

### Potential solution

If uneven cutting occurs, stop the vibratome immediately and carefully return the blade to its resting position. Inspect the blade for visible cracks or other damage; if compromised, replace it with a new one before proceeding. If the blade appears intact, remove the tray to access the blade holder and use the VibroCheck to assess and adjust vibration settings. Ensure the vertical deflection of the blade is within the recommended range (acceptable value: |∑µm| < 0.2). Once the vibratome is properly calibrated, mount a fresh tissue sample and resume sectioning; the slices should now come out smooth and intact. Regular inspection and maintenance of the vibratome setup can help to prevent this issue.

### Problem 2

Downstream signs of RNA degradation, such as low-quality cDNA profiles on the Bioanalyzer, can compromise both data quality and biological interpretation. This issue often arises from prolonged handling times or high temperature during nuclei isolation and processing.

### Potential solution

To protect RNA from degradation, it is essential to minimize the time between thawing and lysis by performing each step quickly yet carefully. Keep all samples and reagents on ice or at 4°C throughout the protocol. Avoid processing more than 15 samples per person in a single run, as larger batches increase handling times and the risk of RNA decay. Regularly wipe down surfaces and tools with RNase inhibitors like RNaseZAP, and ensure that only RNase-free reagents, tips, and consumables are used. Implementing these precautions will help maintain high RNA quality suitable for downstream sequencing analyses.

### Problem 3

A low percentage of human sequencing reads often indicates that too few graft-derived nuclei were recovered, or that human and host nuclei were not properly separated. This reduces the accuracy of downstream analysis and limits the biological conclusions that can be drawn from the graft data.

### Potential solution

Refine graft dissection under a stereomicroscope to ensure thorough isolation of human tissue, and minimize contamination with host cells. If the graft spans multiple brain areas, consider labeling the grafted cells with a fluorescent marker like nuclear GFP, and include a sorting gate for GFP to help isolate nuclei more accurately. For existing datasets, confirm that the sequencing reads are aligned to a combined human (GRCh38) and rat (mRatBN7.2) reference genome using tools like Cell Ranger. This dual-genome alignment is key to correctly identifying human nuclei and ensuring reliable quantification of graft-derived reads.

### Problem 4

Poor nuclear yield or incomplete tissue lysis during nuclei extraction step can lead to cell clumps or large tissue fragments, reducing the number of recoverable nuclei and compromising downstream analyses. This issue often stems from insufficient mechanical dissociation or suboptimal lysis conditions.

### Potential solution

During homogenization, use both the loose and tight pestles, applying 10–15 firm strokes with each to thoroughly break down the tissue. If tissue clumps remain, perform additional strokes using the tight pestle while rotating it, until the lysate appears uniformly cloudy with no large fragments. For mature or denser grafts, slightly increase the lysis buffer volume and allow the tissue to sit in the buffer for 1-2 minutes before douncing. Always keep samples and reagents ice-cold or at 4°C to preserve RNA quality and maintain nuclear integrity.

### Problem 5

Nuclei adhesion to the walls of collection tubes during sorting can cause sample loss and reduce material available for downstream analysis. This typically occurs when collection tubes are not properly coated.

### Potential solution

To reduce nuclei adhesion to the collection tubes, pre-coat the tubes with resuspension buffer at least 30 minutes before sorting. This gives the buffer time to coat the surface and reduce stickiness. Keep the tubes on ice until use. DNA LoBind tubes are strongly recommended, as they minimize sample loss and help preserve nuclei quality during sorting. Before use, ensure any residual liquid is removed from the collection tubes.

### Problem 6

High background during fluorescence-activated nuclei sorting (FANS) or frequent clogging of the sorter can slow down the procedure and compromise data quality. This often results from insufficient debris removal or overly dense nuclear suspensions.

### Potential solution

Optimize the douncing step to effectively release nuclei while minimizing tissue disruption that generates debris. If the lysate appears particularly dense, dilute it with additional cold resuspension buffer prior to sorting to reduce viscosity. When working with cell pellets or organoid samples, filtering through a 40 µm nylon mesh can help remove large debris and prevent sorter clogs. However, it is recommended to avoid this filtration step for graft-derived samples, as it can significantly reduce nuclei yield.

### Problem 7

Low purity of sorted nuclei during FANS can compromise downstream analyses by introducing contaminating debris or unwanted cell populations. This often arises from suboptimal gating parameters or inconsistent nuclear extraction.

### Potential solution

Use freshly prepared Draq7 solution at the recommended 1:1000 dilution to ensure reliable nuclear staining. Before sorting your samples, analyze 200-500 events from the most abundant sample to verify and optimize gating strategy. Include appropriate controls - such as unstained samples and a DNA stain-only control – to clearly define the nuclear population and exclude debris or doublets. Review and adjust the gating strategy with each new run to maximize the quality of your sorted nuclei.

### Problem 8

High variability between samples often results from differences in tissue dissection, nuclei isolation, or processing time. These inconsistencies can introduce batch effects that make it harder to identify real biological differences across your samples.

### Potential solution

Maintain consistency in sample handling by standardizing timing and conditions during both dissection and nuclei extraction. When processing multiple samples, distribute the workload among researchers to minimize unnecessary delays. Ensure all samples are processed at the same temperature and within comparable time windows. During downstream analysis, apply batch correction tools like Harmony^22^ to reduce residual technical variation, enabling more reliable integration of data from different grafts or experiments and improving overall reproducibility.

### Resource availability

- ***Lead contact***: *Further information and requests for resources and reagents should be directed to and will be fulfilled by the lead contact, Prof. Alessandro Fiorenzano*.
- ***Technical contact***: *Technical questions on executing this protocol should be directed to and will be answered by the technical contact, Dr. Edoardo Sozzi*.
- ***Materials availability***: *This study did not generate new unique materials. All materials used in this protocol are purchasable*.
- ***Data and code availability***: *Raw single-nucleus RNA sequencing data generated using this protocol are publicly available at the Gene Expression Omnibus (GEO) under accession number GSE233885, as part of the datasets published in Fiorenzano et al*.^*1*^ *Additional data and analysis code related to this protocol are available from the corresponding authors upon request*.

## Acknowledgments

The authors thank Anna Hammarberg, Jenny G. Johansson, and Michael Sparrenius for their expert technical support and assistance in establishing this methodology. They also thank Prof. Malin Parmar, Andreas Bruzelius and Maria Garcia Garrote for valuable input during protocol development. The authors further acknowledge the FACS platform at MultiPark (Multidisciplinary Research in Parkinson’s Disease), Lund University, for their support with nuclei sorting, and the Center for Translational Genomics (CTG), Lund University, for providing sequencing services. This work has been supported by the Swedish Research Council (A.F. 2022-01432), and the Swedish Parkinson Foundation (A.F. Parkinsonfonden). E.S. was supported by the Royal Physiographic Society in Lund (44435 and 43374).

## Author contributions

Conceptualization, E.S., A.F.; Data acquisition, E.S., A.F.; Data analysis, E.S., P.S.; Writing and editing, E.S., P.S., A.F.

